# Functional loss of CHS2 confers high levels resistance to *Bacillus thuringiensis* Vip3Aa in *Spodoptera exigua* and *Agrotis ipsilon*

**DOI:** 10.1101/2025.06.05.658017

**Authors:** Peng Wang, Zhenxing Liu, Qiangqiang Kang, Chongyu Liao, Luming Zou, Kaikai Mao, Hui Yao, Yiping Li, Yutao Xiao

**Author notes:** These authors contributed equally to this work (Peng Wang, Zhenxing Liu).

## Abstract

*Bacillus thuringiensis* (Bt) crops, engineered to produce insecticidal proteins such as Vip3Aa and Cry toxins, have revolutionized pest management by providing a sustainable alternative to chemical pesticides. However, the development of resistance to these toxins has driven the investigation of underlying mechanisms to better understand how pests evade toxicity and to develop more effective resistance management strategies. Recently, a laboratory-selected *Spodoptera frugiperda* strain exhibited a high level of resistance to Vip3Aa (resistance ratio: 5562-fold), with the mutation of the chitin synthase gene, *SfCHS2*, identified as a key factor. In this study, we extend these findings to additional lepidopteran species, including *Spodoptera exigua* and *Agrotis ipsilon*. Our results show that *CHS2* knockout strains lack the peritrophic matrix (PM), while the resistant Sfru_R3 strain retains its PM. Knockout of this gene resulted in high levels resistance to Vip3Aa. These findings further validate the role of the *CHS2* gene in Vip3Aa resistance and highlight its potential as a target for resistance management in lepidopteran pests. This study advances our understanding of the molecular mechanisms behind resistance to Vip3Aa and supports the development of strategies to delay resistance in Bt crop management.

## 1. Introduction

Genetically modified (GM) plants that produce toxins derived from the bacterium *Bacillus thuringiensis* (Bt) are known as Bt crops and represent one of the most successful applications of genetic engineering in agriculture. These transgenic plants (eg: cotton, maize and soybean) can produce toxic proteins to disrupt the digestive system of many key lepidopteran and coleopteran pests, thus have revolutionized pest management by offering an efficient, environmentally friendly alternative to chemical pesticides. The most widely used are crops expressing Cry toxins, which have been commercialized and widely grown around the world for decades(Hutchison et al., 2010; Lu et al., 2012). However, intensive planting of these Bt crops has created a growing number of resistance in target insect species (Dhurua and Gujar, 2011; Orozco-Restrepo et al., 2024; Tabashnik et al., 2023; Yang et al., 2022a; Zhang et al., 2016). The Vip3Aa toxin, which is produced in the vegetative stage of Bt, was introduced in crops alone, or with other Cry toxins together, to confer higher protection and delay insect resistance (Kurtz et al., 2007; Niu et al., 2021; Rabelo et al., 2020; Yang et al., 2022a).

Understanding the mechanism of Vip3Aa toxicity and resistance is essential for effective pest management (Wu, 2014; Yang et al., 2022b). The protein sequence of Vip3Aa shows no homology with Cry proteins (Estruch et al., 1996), and current evidence suggests no cross-resistance between the two (Bachler et al., 2024; Chakroun et al., 2016; Kurtz et al., 2007; Liu et al., 2024; Sena et al., 2009), indicating a different resistance mechanism for Vip3Aa. Although the resistance to Vip3Aa in field pest populations is rare (Amaral et al., 2020; Yang et al., 2021a), several laboratory-isolated strains have shown medium-to-high resistance, providing valuable resources for detecting the underlying mechanism (Bachler et al., 2024; Bernardi et al., 2016; Carrière et al., 2023; Jin et al., 2023; Kerns et al., 2023; Liu et al., 2024; Pickett et al., 2017; Roy et al., 2025a, b; Yang et al., 2020; Yang et al., 2021b).

Only a few genes have been identified as being associated with Vip3Aa resistance in vivo. In *Helicoverpa armigera, HaVipR1* was identified as a crucial determinant of Vip3Aa resistance from two field-derived resistant lines, with its role in mediating resistance confirmed through knock-out experiments (Bachler et al., 2024). The involvement of this gene in Vip3Aa resistance was further validated in *Spodoptera frugiperda* (Zhang et al., 2024). Recently, two genes associated with Vip3Aa resistance in *S. frugiperda* were identified from two distinct field-derived populations in our group. The downregulation of *SfMyb* gene in DH-R strain confers 206-fold resistance (Jin et al., 2023), while a naturally occurring mutation of a chitin synthase gene (*SfCHS2*) in Sfru_R3 strain confers 5560-fold resistance (Liu et al., 2024).

The *CHS2* gene is primarily expressed in insect gut epithelial cells and produces chitin for the formation of the peritrophic matrix (PM) (Merzendorfer and Zimoch, 2003). However, the relationship between PM formation and Vip3Aa resistance has not been well defined (Jiang et al., 2023). Following the knock-out of *CHS2* genes in *S. frugiperda, Spodoptera litura*, and *Mythimna separata* in our last paper (Liu et al., 2024), we here extend the knock-out to two additional lepidopteran species—*Spodoptera exigua*, and *Agrotis ipsilon*. The results showed high levels resistance to Vip3Aa and the absence of PM in all knock-out strains, while the PM was retained in the Sfru_R3 strain. These findings provide further evidence of *CHS2* gene’s association with Vip3Aa resistance in lepidopteran insects in general.

## 2. Materials and methods

### 2.1. Insect strains and rearing

The *S. frugiperda, S. litura*, and *M. separata* strains were described in our previous work (Liu et al., 2024). The Se-WT strain of *S. exigua* was first collected in Yantai City, Shandong Province, China, in 2023. The Ai-WT strain of *A. ipsilon* was purchased from Keyun Biological Co., Ltd. (China) in 2024. The wild-type strains and their corresponding knock-out strains, SeCHS2-KO and AiCHS2-KO, were maintained at 26 ± 2 °C and 70 ± 10% relative humidity (RH) under a 14 h light:10 h dark photoperiod. The larvae were fed with a wheat germ based diet (Jin et al., 2021). Adults were provided with 10% sucrose solution.

### 2.2. Design and preparation of sgRNAs

The *SeCHS2* and *AiCHS2* genes were identified by using the protein sequence of SfCHS2 (protein_id: WXB24377.1) to perform a BLAST search against the *S. exigua* and *A. ipsilon* data base. The sgRNA target sites of *SeCHS2* and *AiCHS2* were then identified using the sgRNAcas9 (V3.0) software (www.biootools.com/software). Two sgRNAs were used for each insect species (Se-sgRNA1 targeting exon21 of *SeCHS2*: 5’-TGTCCACCACTAAATTGGATTGG-3’; Se-sgRNA2 targeting exon22 of *SeCHS2*: 5’- ACGACTGAACACCGACGACCTGG-3’; Ai-sgRNA1 targeting exon21 of *AiCHS2*: 5’- TCTCTACCACGCATCTCAACTGG-3’; Ai-sgRNA2 targeting exon22 of *AiCHS2*: 5’- ACGTCTGAACACAGACGACTTGG-3’. PAM sequences are underlined). The GeneArt Precision gRNA Synthesis Kit (Thermo Fisher Scientific, cat. no. A29377) was used to synthesize and purify the sgRNAs according to the manufacturer’s instructions.

### 2.3. Embryo collection and microinjection

Freshly laid eggs (within 1 hour after oviposition) were collected and fixed onto a glass slide with double-sided tape. A mixture of sgRNAs (200 ng/μl) and Cas9 protein (500 ng/μl) (TrueCut^TM^ Cas9 Protein v2, Thermo Fisher Scientific, cat. no. A36497) was injected into individual eggs using a FemtoJet 4i and InjectMan 4 microinjection system (Eppendorf, Hamburg, Germany) within 1 hour. Injected eggs were placed at 26 ± 2^°^C with 70 ± 10% relative humidity under a 14 h light:10 h dark photoperiod until hatching.

### 2.4. Mutagenesis detection and homozygous screening

Genomic DNA was extracted from the hind legs of adults prior to mating. Primers flanking the target sites were used to detect mutagenesis (Table S1), and the PCR product was sequenced by Sangon Biotech (Shanghai, China). Microinjected G_0_ individuals were single paired with wild-type individuals to generate G_1_ progeny. Heterozygous G_1_ adults with the same mutation were crossed to generate G_2_ progeny. The homozygous G_2_ individuals were then crossed to establish a knock-out strain.

### 2.5. Bt toxin and bioassays

The Vip3Aa protoxin was purchased from Beijing Generalpest Biotech Research Co., Ltd. (Beijing, China). Diet overlay bioassays were conducted as previously described (Liu et al., 2024). Briefly, diet was dispensed into 24-well plates, and different concentrations of Vip3Aa (0, 200, 400, and 800μg /cm^2^) were evenly applied to the surface and allowed to dry. A single unfed neonate was transferred to each well. After 7 days, mortality was recorded (larvae were considered dead if they had died or remained in the first or second instar).

SPSS18.0 was used to calculate the LC_50_. LC_50_ values with non-overlapping 95% confidence intervals (CIs) were considered significantly different. Resistance ratios were calculated by dividing the LC_50_ of the tested strain by the LC_50_ of the wild-type or susceptible strain (SS).

For homozygous knock-out individuals that suffered severe fitness cost and could not reach the adult stage, progeny from heterozygous crosses were used for the overlay bioassay. Survivors were subjected to PCR testing to determine their mutagenesis types.

### 2.6. Dissection and histological analysis

The final instar larvae were used for dissection in all strains. The midgut was fixed with Bouin’s solution to observe the presence of PM. For histological analysis of the midgut, the samples were fixed in 4% paraformaldehyde and sent to Servicebio (Wuhan, China) for paraffin embedding. Briefly, the samples were dehydrated using a serial grades of ethanol, followed by clearing with xylene. The pretreated midgut was embedded in paraffin (Sigma-Aldrich, USA), and the midgut samples were cross-sectioned into 5 μm thick slices. The paraffin sections were then deparaffinized through successive baths of xylene and rehydrated through serial grades of ethanol, stained with hematoxylin–eosin (Servicebio, Wuhan, China), and finally mounted with neutral resin (Servicebio, Wuhan, China). The slides were examined under a digital pathology system (Pannoramic MIDI, 3DHISTECH Int.) to assess the morphological characteristics of the midgut samples.

## 3. Results

### 3.1. Knockout of *CHS2* resulted in high levels resistance to Vip3Aa toxin in *S. exigua* and *A. ipsilon*

The *CHS2* gene sequences of *S. exigua* (GenBank ID: DQ912929.1) and *A. ipsilon* (GenBank ID: OL826754.1) were used to design sgRNAs to induce mutations at the target sites. Two strains with frameshift mutations were obtained and designated as SeCHS2-KO and AiCHS2-KO (Fig. S1). Each strain carries a deletion of 5 or 146 base pairs, respectively. The altered sequence of SeCHS2 introduced a stop codon in exon 21, yielding a truncated protein consisting of 1350 amino acids. Similarly, the modified *AiCHS2* sequence introduced a stop codon in exon 23, producing a truncated protein of 1,396 amino acids.

The bioassay results showed that SeCHS2-KO strain survived well on a diet overlaid with 800 μg/cm^2^ Vip3Aa toxin, exhibiting a resistance ratio of >33,333-fold compared to the wild-type susceptible strain (Table 1). However, the homozygous mutants of *A. ipsilon* experienced a very high fitness cost, with only a few individuals reaching the adult stage and no offspring obtained. Alternatively, we used the progeny of heterozygous crosses in the bioassay. Survivors were observed in all treatments, with 13 out of 72 progressing beyond the second instar when exposed to ≥200 μg/cm^2^ (maximum: 800 μg/cm^2^) Vip3Aa toxin (Table 2). PCR analysis showed that all these survivors were homozygotes (Fig. S2). The proportion of survivors did not significantly differ from the expected 25% homozygous ratio, as determined by a one-sample z-test for proportions (*p* = 0.174). These results suggest that the resistance ratio of homozygous mutants of *A. ipsilon* is >11,268-fold compared to the wild-type susceptible strain.

**Table 1.** Responses of CHS2-knockout strains to Vip3Aa.

| Strain | n<sup>a</sup> | LC<sub>50</sub> (95% CI)<sup>b</sup> | Slope ± SE | RR<sup>c</sup> |
| --- | --- | --- | --- | --- |
| Sf-SS<sup>d</sup> | 192 | 0.13 (0.11 - 0.16) | 4.0 ± 0.6 | 1.0 |
| SfCHS2-KO-A<sup>d</sup> | 80 | > 1600<sup>d</sup> | NA | > 12,000 |
| SI-SS<sup>d</sup> | 96 | 0.016 (0.0030 - 0.086) | 4.1 ± 0.8 | 1.0 |
| SICHS2-KO<sup>d</sup> | 96 | >1600<sup>d</sup> | NA | > 100,000 |
| Ms-SS<sup>d</sup> | 168 | 1.20 (0.97 - 1.48) | 6.6 ± 1.0 | 1.0 |
| MsCHS2-KO<sup>d</sup> | 96 | >1600<sup>d</sup> | NA | > 1,300 |
| Se-SS | 192 | 0.024(0.016 - 0.034) | 4.13 ± 0.78 | 1.0 |
| SeCHS2-KO | 96 | >800 | NA | >33,333 |
| Ai-SS | 144 | 0.071(0.032 - 0.10) | 3.68 ± 0.98 | 1.0 |

<sup>a</sup> Number of neonates tested.
<sup>b</sup> Median lethal concentration (LC<sub>50</sub>); concentration that caused 50% of neonates to die or fail to advance to the third instar in 7 days and its 95% CI in µg Vip3Aa per cm<sup>2</sup> diet.
<sup>c</sup> Resistance ratio: LC<sub>50</sub> for a strain divided by the LC<sub>50</sub> for SS.
<sup>d</sup> Data from (Liu et al., 2024).
CI, confidence interval; SS, susceptible strain.

**Table 2.** Bioassay with offspring from heterozygous crosses of *Agrotis ipsilon*.

| Vip3Aa concentration<br>($\mu\text{g}/\text{cm}^2$) | Initial number | Survived number |
| --- | --- | --- |
| 0 | 24 | 23 |
| 200 | 24 | 6 |
| 400 | 24 | 3 |
| 800 | 24 | 4 |

### 3.2. The peritrophic matrix was absent in the *CHS2* knockout strains

Since CHS2 is responsible for chitin synthesis in the midgut of lepidopterans, it’s important to determine whether knockout of this gene affects midgut structure. Knockout strains and their corresponding wild-type strains were dissected at their final instar of larval stage. The midgut tissues were excised and fixed in Bouin’s solution, which imparts a bright yellow coloration to enhance visibility. Because the midgut and PM have different contraction ratios, the PM protrudes from the midgut when excised, providing a clear indicator of its presence. The results showed the PM were absent in all knockout strains but present in wild-type strains (Fig. 1).

**Fig. 1.**
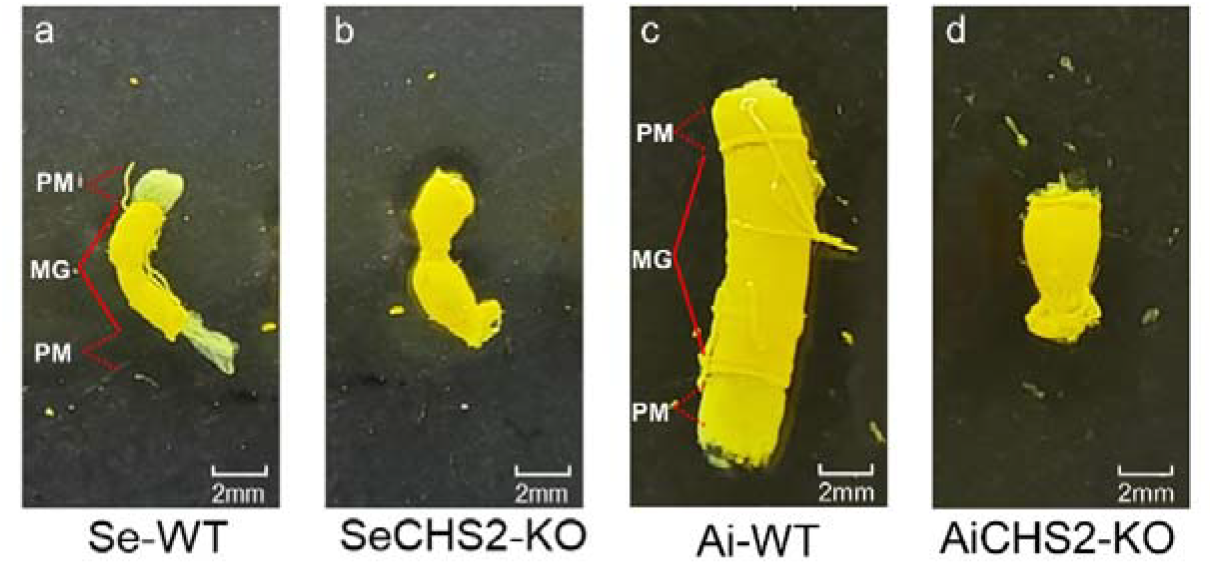
The peritrophic matrix was absent in the CHS2 knockout strains. Dissection of final instar larvae clearly show the presence of the PM in wild-type strains of S. exigua (a) and A. ipsilon (c), but it is absent in their corresponding knockout strains (b and d). MG, midgut; PM, peritrophic matrix.

The CHS2 gene was initially associated with Vip3Aa resistance in a laboratory-selected S. frugiperda strain (Sfru_R3). To further investigate this, we also dissected the Sfru_R3 strain along with the S. frugiperda knockout strain (SfCHS2-KO-A), and the M. separata (MsCHS2-KO) and S. litura (SlCHS2-KO) knockout strains described in our previous study (Liu et al., 2024). The results confirmed that the PM was absent in all knockout strains, but present in Sfru_R3 and the corresponding wild-type strains (Fig. S3).

### 3.3. Histological structure analysis further confirmed the absence of PM in knockout strains of two species

The *A. ipsilon* strains were selected for histological structure analysis. Transverse sections of the midgut showed that the PM was absent in the knockout strain AiCHS2-KO, but present in wild-type susceptible strains (Fig. 2).

**Fig. 2.**
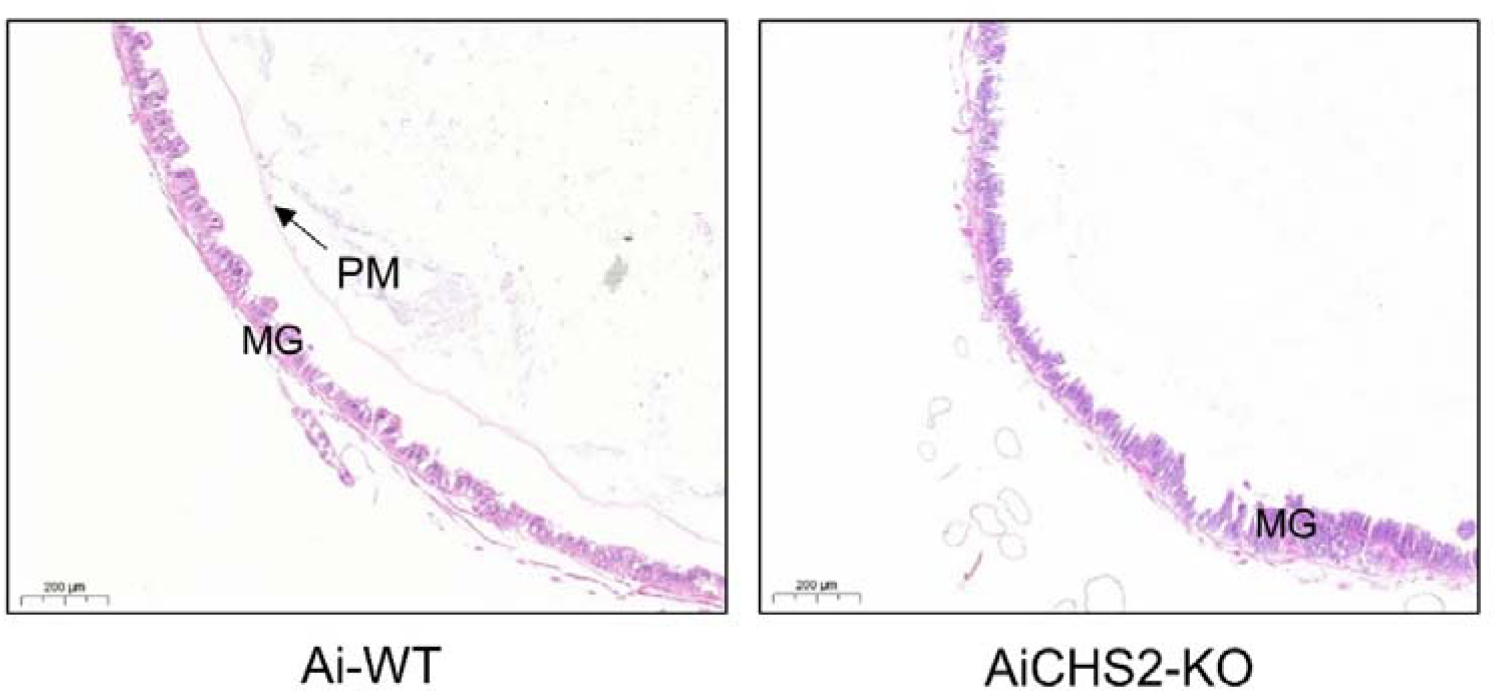
Histological analysis of transverse midgut sections in *A. ipsilon*. The results show that the PM is absent in the *A. ipsilon* knockout strain (AiCHS2-KO), but present in the wild-type strain (Ai-WT). MG, midgut; PM, peritrophic matrix.

## 4. Discussion

In our previous work, we identified a retrotransposon (Yaoer)-mediated disruption of *CHS2* in *S. frugiperda*, which conferred high resistance to Vip3Aa (Liu et al., 2024). Knockout of this gene in *S. frugiperda*, as well as in two other lepidopteran species, *S. litura* and *M. separata*, resulted very high resistance to Vip3Aa. In this study, we generated knockout strains for two additional species, *S. exigua* and *A. ipsilon*, both of which also exhibited high resistance to Vip3Aa (Table 1 and Table 2). The role of CHS2 in determining Vip3Aa susceptibility across five species from three different noctuid tribes suggests a conserved function in Lepidoptera.

The outcomes of CHS2 knockout, however, differed markedly across species. In *A. ipsilon*, only a few homozygous mutants survived to adulthood, and none produced viable offspring. Similarly, our attempts to establish a knockout strain of *H. armigera* failed due to rapid colony collapse. By contrast, a homozygous knockout strain of *S. exigua* could be obtained, although many individuals died during larval development (data not shown), indicating a significant but less absolute fitness cost. Together with our previous findings in three other species(Liu et al., 2024), these results demonstrate that CHS2 disruption severely compromises survival and reproduction, but the magnitude of this effect varies among species. Comparing these species-level outcomes provides critical insight into the likelihood of CHS2-associated resistance persisting in natural populations.

The PM in insects functions as a permeable barrier between the food bolus and the midgut epithelium, facilitating digestive processes while protecting the brush border from mechanical damage and attacks by toxins and pathogens (Tellam, 1996). The PM is composed of a complex mixture of chitin, proteins, glycoproteins and proteoglycans (Wang and Granados, 2001). Our study demonstrates that knocking out the *CHS2* gene resulted in the absence of the PM in all five species during the larval stage, as shown in both dissection results (Fig 1 and Fig S3) and histological structure analysis (Fig 2). Many homozygous individuals died during the larval stage, suggesting impaired digestive capability.

The PM in the Sfru_R3 strain, on the other hand, remained intact (Fig 1), likely due to the reduced yet persistent presence of wild-type *SfCHS2* transcripts (Liu et al., 2024). It appears that the level of wild-type *SfCHS2* transcripts is negative correlated with the resistance ratio to Vip3Aa and, consequently, with the chitin content in the PM. Recent research has shown that the binding of Domain V of Vip3Aa toxin to the PM via glycan-binding activity contributes to Vip3Aa’s insecticidal activity, while domains II-III of Vip3Aa were involved in binding to brush border membrane vesicles (BBMV) proteins (Infante et al., 2024; Jiang et al., 2023).

Understanding how Vip3Aa interacts with its receptor and functions as an efficient insecticide is crucial for countering resistance in the field. It has been proposed that proteins such as scavenger receptor-C (Sf-SR-C), fibroblast growth factor receptor-like protein (Sf-FGFR), and Prohibitin II serve as Vip3Aa receptors in Sf9 cell line (An et al., 2022; Jiang et al., 2018a; Jiang et al., 2018b). However, knockout experiments using CRISPR/Cas9 in *S. frugiperda* failed to confirm Sf-SR-C and Sf-FGFR as functional receptors (Shan et al., 2022). To date, three genes have been reported as being associated with Vip3Aa resistance in *S. frugiperda*: *SfMyb, SfCHS2* and *SfVipR1* (Jin et al., 2023; Liu et al., 2024; Zhang et al., 2024). Additionally, a recent study indicates that three distinct genetic loci are involved in Vip3Aa resistance in five resistant *H. zea* strains (Yang et al., 2024). Collectively, these findings indicate that resistance to Vip3Aa is likely determined by multiple factors, with mutations in *CHS2* having greater potential to confer high levels of resistance.

In summary, our findings demonstrate here that knocking out *CHS2* in multiple lepidopteran species confers high levels resistance to Vip3Aa toxin, with the phenotype closely linked to the absence of the PM. These results highlight the essential role of *CHS2* in maintaining PM integrity and its association with Vip3Aa susceptibility. Further investigation into the involvement of the PM in Vip3Aa resistance presents an intriguing research avenue. A deeper understanding of this mechanism could provide valuable insights into the mode of action of Vip3Aa and contribute to the development of innovative strategies for sustainable pest management.

## Data availability

All data generated and analyzed in this study are available within the article and the supplementary information.

## Acknowledgements

The authors acknowledge the financial support by Biological Breeding-National Science and Technology Major Project (2022ZD04021), theAgricultural Science and Technology Innovation Program (CAAS-ZDRW202412), Innovation Program of Chinese Academy of Agricultural Sciences (CAAS-CSCB-202303), Shenzhen Science and Technology Program (KQTD20180411143628272), the National Natural Science Foundation of China (32202352), and The Agricultural Science and Technology Innovation Program of Chinese Academy of Agricultural Sciences.

