## Supplementary data for "Functional loss of CHS2 confers high levels resistance to *Bacillus thuringiensis* Vip3Aa in *Spodoptera exigua* and *Agrotis ipsilon*"

**Appendix A. Supplementary data**

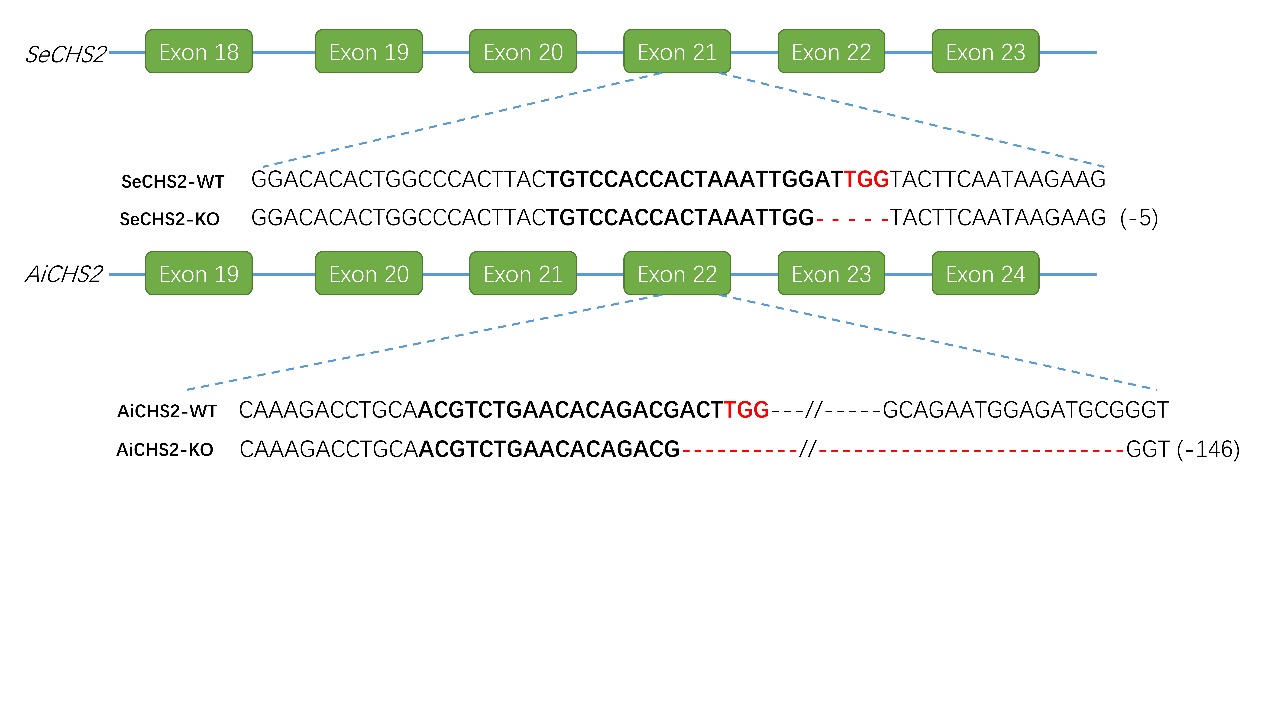

Fig. S1. Schematic diagram of CRISPR/Cas9-mediated mutagenesis of *CHS2* gene in Spodoptera exigua and Agrotis ipsilon. The mutation in *SeCHS2* occurred in exon 21, while the mutation in *AiCHS2* occurred in exon 22. The protospacer adjacent motif (PAM site, 5'-NGG-3') within the blacked sgRNA target sequence is highlighted in red. Compared to the wild-type (WT) sequence, red dashes indicate deletions, with the number of deleted bases shown in brackets.

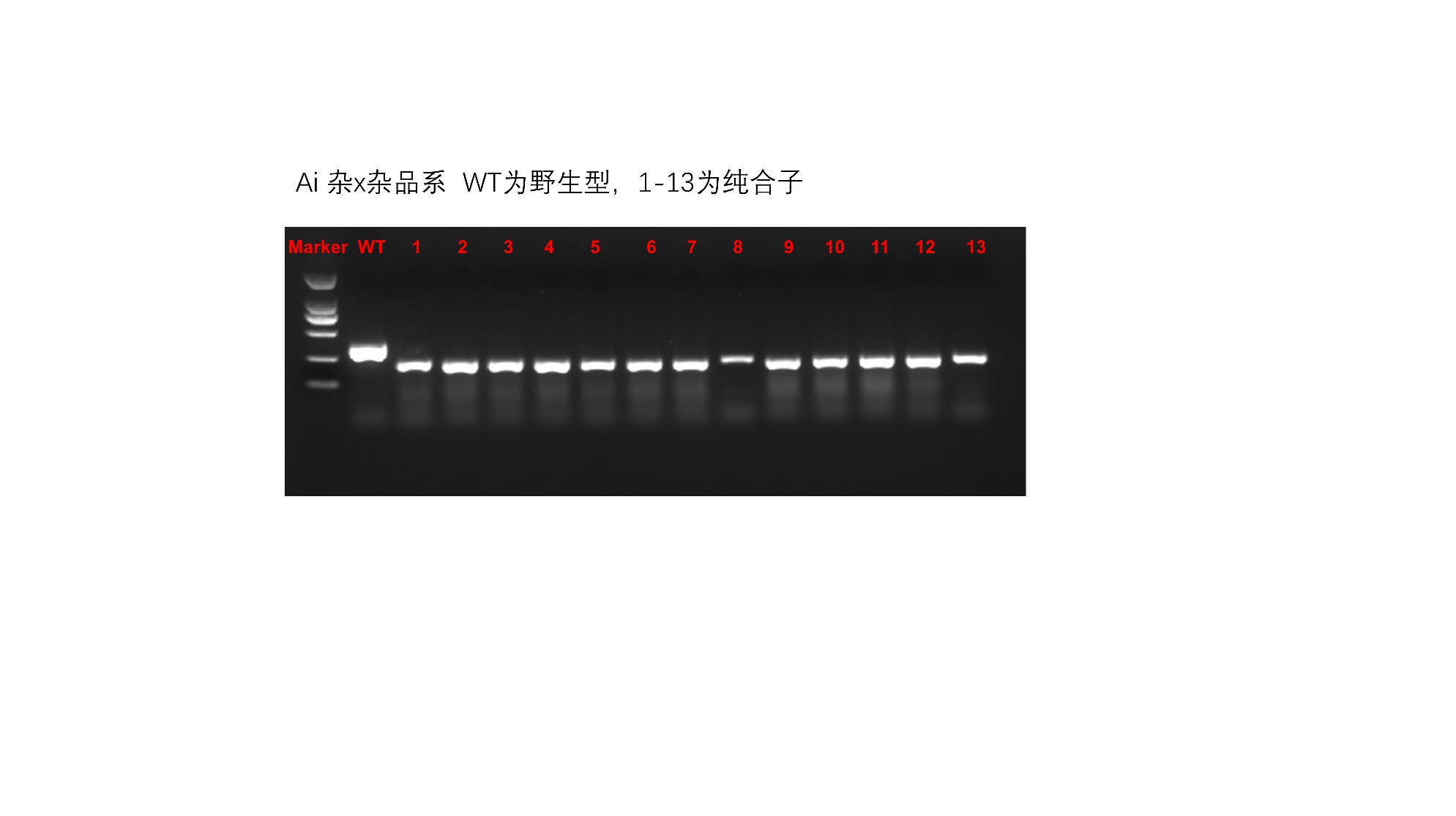

Fig. S2. PCR test for Agrotis ipsilon larvae survived on diet overlay assay. 1-13 represent 13 survived individuals. WT represent a wild type sample. This result indicates all survivals were homozygotes.

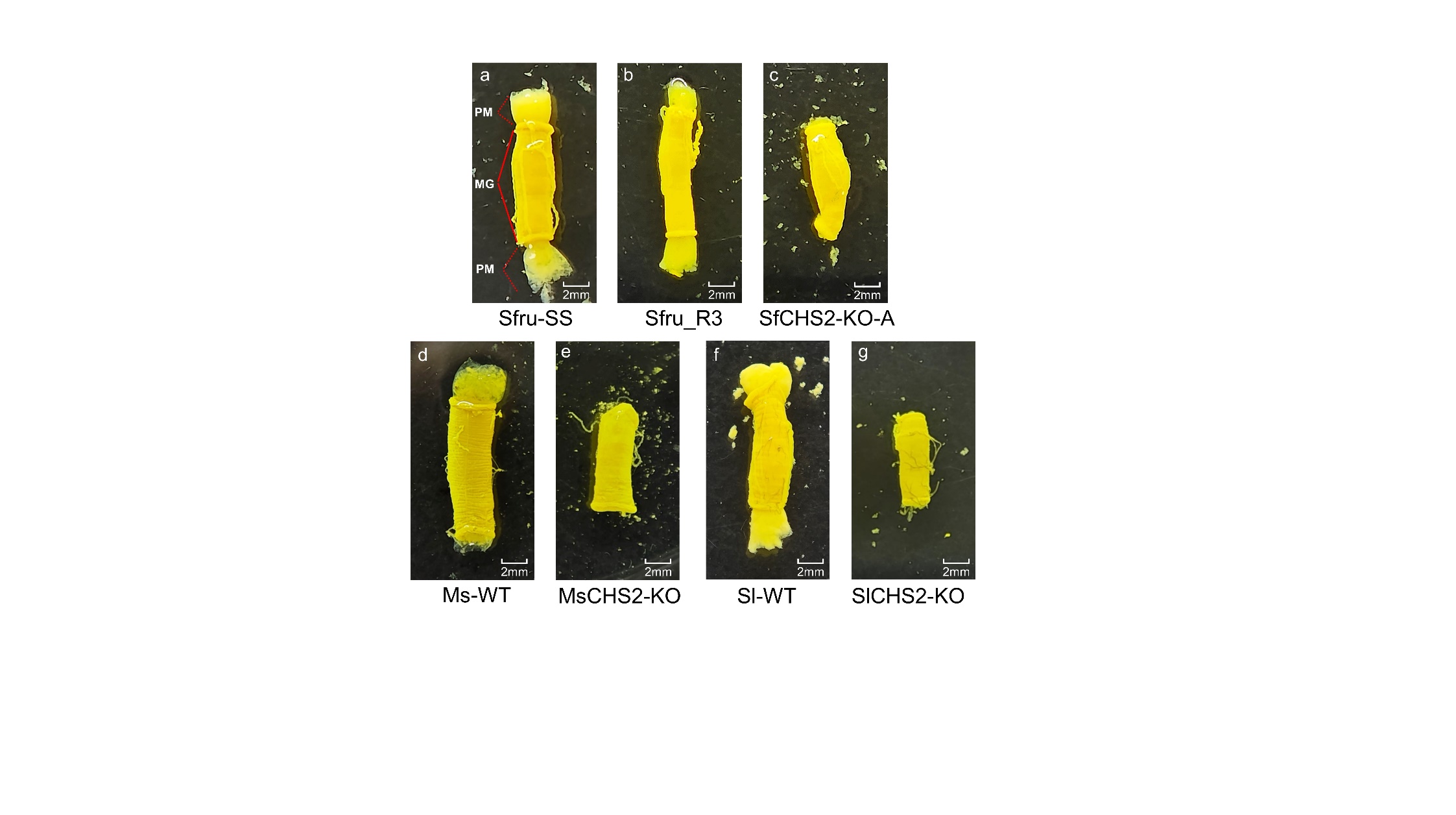

Fig S3. The peritrophic matrix is absent in all the CHS2 knockout strains. Dissection of final instar larvae clearly showed the presence of the PM in *Spodoptera frugiperda* resistance strain (b) and wild-type strains of *S. frugiperda* (a), *Mythimna separate* (d) and *Spodoptera litura* (f), but it was absent in their corresponding knockout strains (c, e and g). MG, midgut; PM, peritrophic matrix.

Table S1 Primer sequences used to detect CRISPR/Cas9 mediated mutation

| Primer ID | Sequence (5’-3’) |
| --- | --- |
| Ai_chs2_F | cgcgcttatagagcagaacg |
| Ai_chs2_R | gatatatccgactccggcgt |
| Se_chs2_F | gttcctcatcttcttcggttcc |
| Se_chs2_R | caagttatgaatggtcttcctcct |
